# Differentiation and Expansion of Tumor-Infiltrating T Cell Clonotypes Occurs in the Spleen Following Immune Checkpoint Blockade

**DOI:** 10.1101/2023.09.04.555982

**Authors:** Duncan M. Morgan, Brendan L. Horton, Maria Zagorulya, J. Christopher Love, Stefani Spranger

**Affiliations:** Department of Chemical Engineering at MIT; Koch Institute at MIT; Department of Biology, MIT

**Author notes:** Authors Contributed Equally (Duncan M. Morgan, Brendan L. Horton).

## Abstract

Immune checkpoint blockade (ICB) enhances tumor-reactive T cell responses against cancer, leading to long-term tumor control and survival in a fraction of patients. Given the increasingly recognized complexity of T cell differentiation that occurs in response to chronic antigen stimulation, it remains unclear precisely which T cell differentiation states are critical for the response to ICB, as well as the anatomic sites at which ICB-mediated reinvigoration of these T cells occurs. We used paired single-cell RNA and T cell receptor (TCR) sequencing to profile endogenous, tumor-reactive CD8^+^ T cells isolated from tumors, tumor-draining lymph nodes, and spleens of mice treated with ICB. We identified an intermediate-exhausted population of T cells in the spleen which underwent the greatest expansion in response to ICB and gave rise to the majority of tumor-infiltrating clonotypes. Increasing concentrations of antigen in the spleen perturbed the differentiation of this phenotype towards a divergent exhausted_KLR state, resulting in reduced numbers of tumor-infiltrating T cells and blunted ICB efficacy. Likewise, an analogous population of exhausted_KLR CD8^+^ T cells in matched human tumor and blood samples and exhibited diminished tumor-trafficking ability. These data demonstrate that the spleen is a critical anatomic site for coordinating the differentiation of tumor-infiltrating clonotypes and their expansion in response to ICB.

## Introduction

Immune checkpoint blockade (ICB) using antagonist antibodies targeting CTLA-4 or PD-(L)1 can provide durable responses against cancer by enhancing CD8^+^ T cell immunity (1–3), for a subset of patients (4). Understanding the mechanisms of response and sources of resistance to ICB could improve the selection of patients likely to respond and aid the rational design of new therapies. Most evidence suggests that ICB works by enhancing the functionality of tumor-reactive CD8^+^ T cells (1–3). Generating an anti-tumor CD8^+^ T cell response is a sequential, multi-step process involving the coordination of a variety of cell types across multiple anatomic locations (5). This process initiates with the priming of naïve CD8^+^ T cells by antigen-presenting cells (APCs) in the tumor-draining lymph node (TdLN) (6, 7). Guided by signals received during priming, tumor-reactive CD8^+^ T cells then undergo clonal expansion and begin to differentiate into effector T cells (8, 9). Tumor-reactive CD8^+^ T cells migrate to the tumor, where they eliminate tumor cells in an antigen-specific manner (10). Clinically-detectable tumors use a variety of potential immune-suppressive mechanisms to avoid engagement from cytotoxic T cells (11), including the chronic expression of tumor antigens, which induces exhaustion among tumor-reactive CD8^+^ T cells (12, 13). This CD8^+^ T cell exhaustion often manifests an increased expression of immune inhibitory checkpoint receptors, including PD-1, CTLA-4, and LAG-3 (14). Engagement of these receptors reduces T cell functionality, but blocking these receptors with ICB can make the T cells more responsive toother costimulatory signals (15, 16), promote clonal expansion of functional CD8^+^ effector cells (17), and enhance migration of CD8^+^ effector cells to the tumor. Together, these factors can reinvigorate an anti-tumor immune response.

Studies in chronic viral infection and cancer have demonstrated that T cell exhaustion represents a distinct trajectory of differentiation that extends beyond terminally exhausted T cells (18–22). An exhausted CD8^+^ T cell response comprises heterogenous differentiation states, including TCF-1^+^ “stem-like” progenitor exhausted T cells that maintain an exhausted response through constant differentiation and self-renewal (18–22), and exhibit an enhanced ability to response to ICB (23, 24). Both mouse cancer models and human tumors have shown TCF-1^+^ CD8^+^ T cells are critical for ICB responses, but the anatomic locations from which these ICB-responsive CD8^+^ T cells emerge is unclear (25–29). Both TdLNs and tumors have been proposed as sites that maintain ICB-responsive CD8^+^ T cell populations (25–29), but peripheral expansion also appears critical for generating reinvigorated tumor-infiltrating CD8^+^ T cells (30–32). Recent single-cell analyses of antigen-specific CD8^+^ T cells in LCMV-infected mice have highlighted additional transcriptional phenotypes associated with exhaustion, including intermediate exhausted populations that can give rise to multiple distinct terminally exhausted fates (33, 34). Nonetheless, the precise transcriptional states of T cells providing anti-tumor immunity, the differential contributions of the distinct phenotypes of T cells providing tumor control in response to ICB, and the anatomic locations from which responses to ICB emerge remain unclear.

Here, we used paired single-cell RNA and TCR sequencing to profile the endogenous, anti-tumor T cells isolated from the tumors, TdLNs, and spleens of mice treated with ICB. We found high concordance between the transcriptional states present among these tumor-specific T cells and the transcriptional states previously identified in chronic viral infection (33, 34), but we also found clear biases for distinct differentiation states across anatomic locations. Surprisingly, the spleen was a critical anatomic site for coordinating the differentiation of an “intermediate-exhausted” CD8^+^ T cell population into either a terminally-exhausted phenotype comprising the majority of tumor-infiltrating cells, or into an exhausted_KLR phenotype that exhibited minimal infiltration into the tumor. Using adoptive transfer assays with TCR transgenic T cells, we confirmed the ability of this splenic intermediate-exhausted population to expand in response to ICB and seed other anatomic sites, including the tumor and TdLN. Introducing antigen into the spleen skewed the differentiation of this intermediate population towards the exhausted_KLR phenotype, leading to reduced numbers of tumor-infiltrating T cells and blunted responses to ICB. Using published data sets, we identified corroborating evidence for human T cells manifesting this exhausted_KLR transcriptional state and determined that this differentiation state demonstrated limited infiltration into tumors, despite clonal expansion. These results demonstrate that the spleen is a critical anatomic site for the expansion of tumor-infiltrating clonotypes in response to ICB and that perturbing the differentiation of these clonotypes to favor the exhausted_KLR state can blunt tumor infiltration and efficacy of ICB.

## Results

### SIY-reactive CD8^+^ T cells accumulate in the Spleen during ICB

To demonstrate the impact of ICB on tumor growth, we inoculated mice subcutaneously with KP tumor cells and treated mice with ICB (Figure 1A). Consistent with our previous report (9), ICB delayed the growth of subcutaneous KP tumors (Figure 1B). To track tumor-reactive CD8^+^ T cells during ICB, we used a KP cell line engineered to express the model antigen SIY, which binds H2-K^b^ (KP.SIY). We inoculated mice subcutaneously with KP.SIY and treated mice on days 7 and 10 following tumor inoculation with ICB (Figure 1C). KP.SIY tumors treated with ICB showed reduced tumor growth at day 14 post tumor inoculation (Figure 1D). On day 14, we quantified the number of SIY-reactive CD8^+^ T cells in the TdLNs, spleens, and tumors of mice (Figure 1E). We observed an increased absolute number and frequency of SIY-reactive CD8^+^ T cells in the spleens of mice at day 14 post tumor inoculation (Figure 1E, 1F). To determine whether the increase in SIY-reactive CD8^+^ T cells resulted from an accumulation of circulating SIY-reactive T cells, rather than expansion of spleen-resident populations, we injected tumor-bearing mice intravenously with fluorescently-labeled anti-CD45 antibody before euthanasia to label blood-accessible cells. ICB-treated mice had a lower percentage of blood-accessible splenic SIY-reactive CD8^+^ T cells (Figure 1F), indicating that the observed increase in numbers resulted primarily expansion within the spleen itself.

**Figure 1:**
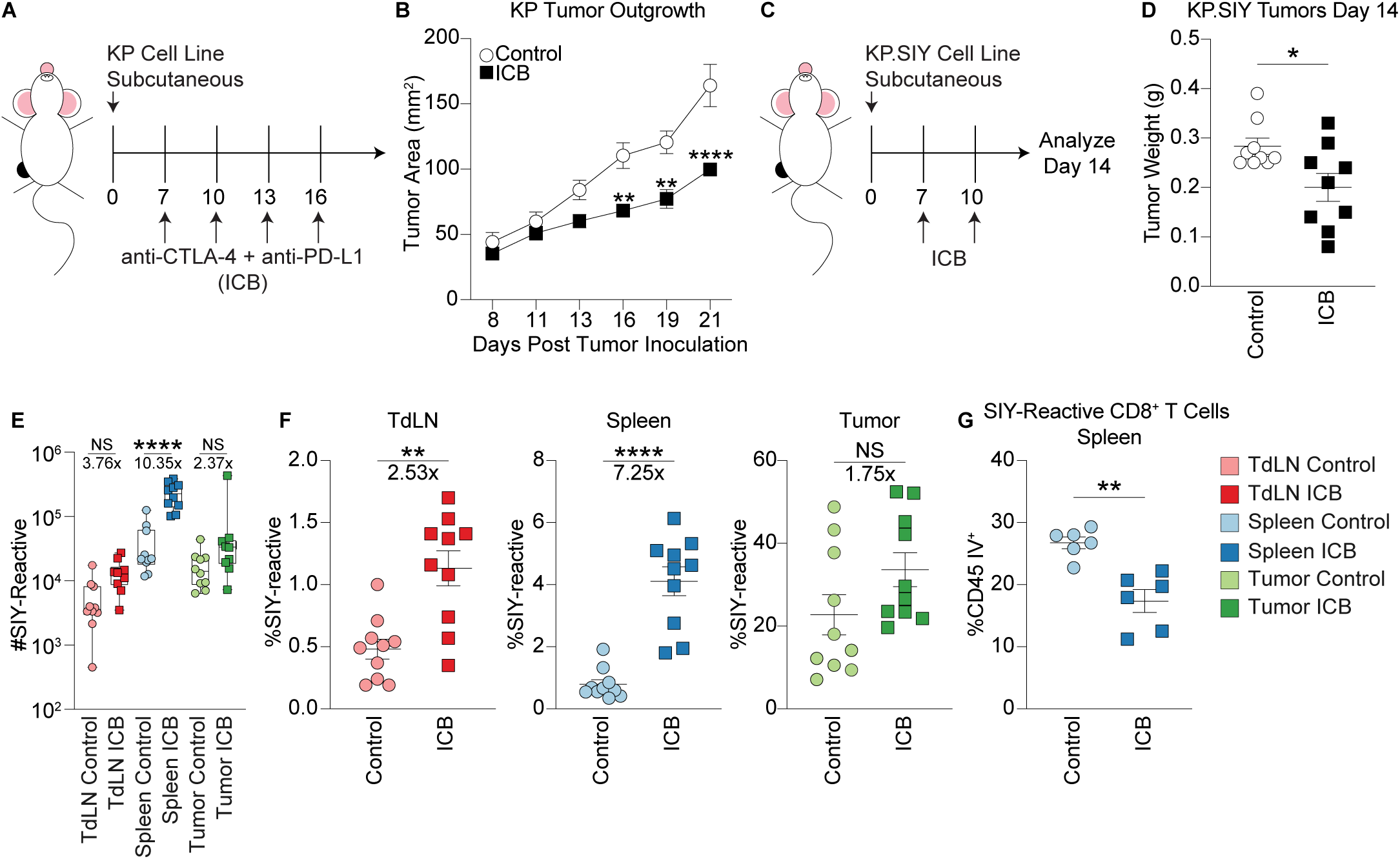
Tumor-Reactive CD8+ T Cells Accumulate in Spleens of ICB-Treated Mice. A) ICB treatment scheme of mice bearing subcutaneous KP tumors. B) KP flank tumor outgrowth, n=6, p-values calculated using 2-way ANOVA. C) ICB treatment scheme of mice bearing subcutaneous KP.SIY tumors. D) Weight of day 14 KP.SIY tumors, n=9, p-values calculated with a Mann-Whitney U test. E) Number of SIY-reactive CD8+ T cells in TdLN, spleen, and tumor on day 14. Fold changes calculated using the median value from each condition, n=10, p-values calculated with one-way ANOVA. F) Percent of CD8+ T cells that are SIY-reactive in TdLN, spleen, and tumor. Fold changes calculated using the median value from each condition, n=10, p-values calculated with a Mann-Whitney U test. G) Percentage of blood-accessible splenic SIY- reactive CD8+ T cells labeled by intravenous inoculation of a fluorescent anti-CD45 antibody administered 3 minutes before euthanasia, n=6. P-values calculated with a Mann-Whitney U test.

### Transcriptional states present among SIY-reactive CD8**^+^** T cells comprise the full spectrum of T cell exhaustion

To determine what phenotypic and clonotypic features underlie the distinct expansion potential of SIY-reactive T cells in the spleen, we then sorted endogenous SIY-reactive T cells from the tumor, TdLN, and spleen of five KP tumor-bearing mice undergoing no treatment (control) and five KP tumor-beading mice undergoing ICB treatment. We analyzed isolated tumor-reactive T cells with single-cell RNA and TCR sequencing (Figure 2A). We performed uniform manifold approximation and projection (UMAP) for dimensionality reduction and identified 10 distinct transcriptional states using unsupervised Louvain clustering (Figure 2B, 2C).

**Figure 2:**
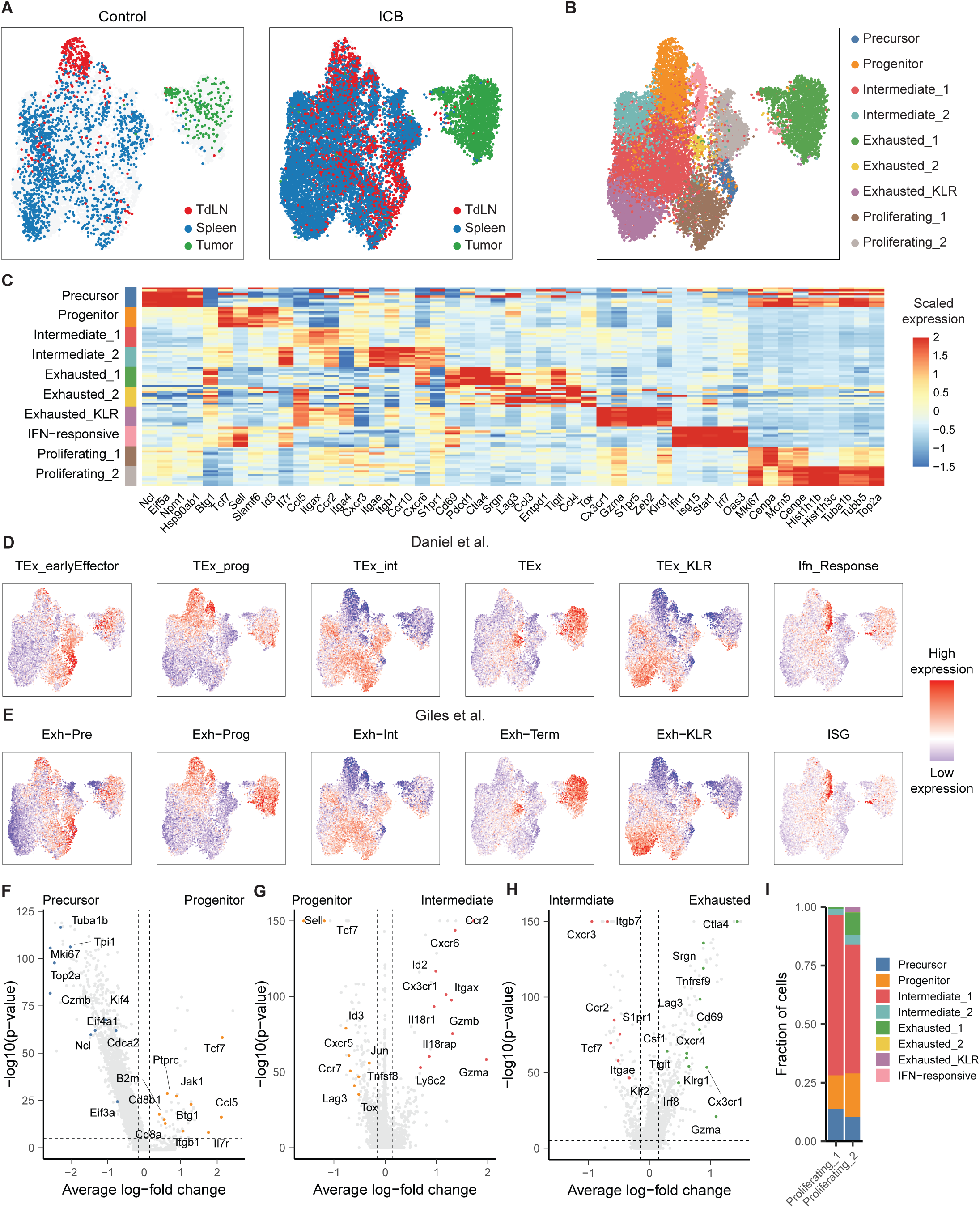
Single-cell, multi-tissue atlas of SIY-reactive CD8+ T cells. A) UMAP of SIY-reactive CD8+ T cells from untreated or ICB-treated KP tumor-bearing mice recovered colored by tissue. B) UMAP of SIY-reactive CD8+ T cells colored by phenotype. C) Heat map of scaled gene expression of marker genes in SIY-reactive CD8+ T cell phenotypes. Each row represents cells from one phenotype recovered from an individual mouse. D) UMAP of gene expression signatures defined among splenic gp33-reactive CD8+ T cells recovered from day 21 of clone 13 LCMV infection by Daniel et al. E) UMAP of gene expression signatures defined among splenic gp33-reactive CD8+ T cells recovered from days 15 and 30 of clone 13 LCMV infection by Giles et al. F) Volcano plot of transcripts differentially expressed between pre-exhausted cells and progenitor-exhausted cells. G) Volcano plot of transcripts differentially expressed between intermediate exhausted cells (Intermediate_1 and Intermediate_2) and progenitor exhausted cells. H) Volcano plot of transcripts es differentially expressed between terminally exhausted cells (Exhausted_1, Exhausted_2, and Exhausted_KLR) and intermediate exhausted cells (Intermediate_1, Intermediate_2). I) Label transfer of non-proliferating cell phenotypes onto proliferating phenotypes. P-values for volcano plots are calculated using a two-sided Wilcoxon rank-sum test and are adjusted using Bonferroni correction.

To identify where these transcriptional states projected on a trajectory of T cell exhaustion, we generated gene expression signatures using two single-cell atlases of CD8^+^ T cell transcriptional phenotypes in LCMV infection (33, 34) and overlaid the phenotypic signatures onto our data. (Figure 2D, 2E). Based on these signatures, we annotated our data with one cluster of precursor-exhausted cells, one cluster of progenitor-exhausted cells, two clusters of intermediate-exhausted cells, two cluster of terminally-exhausted cells, one cluster of exhausted-KLR cells, one cluster that upregulated transcripts associated with interferon-response, and two clusters of proliferating cells (Figure 2B, 2C). Consistent with these annotations, both precursor and progenitor T cells expressed high levels of the transcript *Slamf6* (Figure 2C). In addition, precursor-exhausted T cells were highly proliferative, upregulated transcripts associated with translation and protein synthesis, and downregulated the transcript *Btg1*, while progenitor-exhausted T cells expressed higher levels of the transcription factor *Tcf7* (Figure 2C, 2F). Relative to progenitor-exhausted T cells, intermediate-exhausted T cells expressed elevated levels of transcripts associated with tissue homing (*Ccr2*, *Cxcr6*, *Itgax*) and effector function (*Gzma*, *Gzmb*) and downregulated *Sell* and *Tcf7* (Figure 2C, 2G). Among these two states of intermediate-exhausted cells, intermediate_2 upregulated transcripts such as *Itgae* and *Ccr10*, suggesting that they might comprise precursors of tissue-resident memory T cells (35, 36), while the intermediate_1 upregulated transcripts such *Itga4*, *Gzmk*, and *Eomes*, consistent with a tissue-homing effector T cells (Supplemental Figure 1A). Lastly, compared to both intermediate-exhausted clusters, terminally-exhausted T cells upregulated transcripts such as *Lag3*, *Tnfrsf9*, and *Cd69* (Figure 2C, 2H). Among terminally-exhausted T cells, exhausted_1 was distinguished by the expression high levels of transcripts associated with cytokine signaling (*Ifngr1*, *Il12rb2*), while exhausted_KLR was differentiated by the upregulation of transcripts encoding killer cell lectin-like receptors (KLRs) (*Klrg1, Klrk1*) as well as *Cx3cr1* (Supplemental Figure 1B-D). Lastly, interferon-responsive cells upregulated a variety of transcripts associated with interferon sensing (Supplemental Figure 1E).

To annotate the remaining two states of proliferating cells according to their progression towards terminal T cell exhaustion, we performed label transfer of cluster identities from nonproliferating to proliferating T cells (Figure 2I). These results demonstrated that the majority of proliferating T cells had transcriptional states similar to either a *Slamf6*^+^ progenitor-exhausted phenotype or a *Cxcr3*^+^ intermediate_1 phenotype. In sum, these results demonstrate the transcriptional phenotypes present in this tumor model recapitulate the full spectrum of transcriptional phenotypes that have been observed in models of chronic viral infection in mice. From this point forward, we refer to these transcriptional states as: precursor, progenitor, intermediate_1, intermediate_2, exhausted_1, exhausted_2, exhausted_KLR, interferon-responsive, proliferating_1, and proliferating_2.

### Transcriptional states associated with exhaustion vary in frequency between the tumor, lymph node, and spleen

We next examined the frequency of each transcriptional phenotype among the recovered SIY-reactive T cells from the tumor, TdLN, and spleen (Figure 3A, Supplemental Figure 2). We found that in both control and ICB-treated mice, the TdLN was enriched in transcriptional states associated with early stages in the progression towards exhaustion, including precursor T cells and progenitor T cells. In contrast, the spleen contained elevated frequencies of intermediate exhausted T cell states, and the tumor-infiltrating T cells almost entirely comprised terminally exhausted phenotypes. These data suggest that these tissues are populated by increasingly differentiated phenotypes of tumor-reactive T cells.

**Figure 3:**
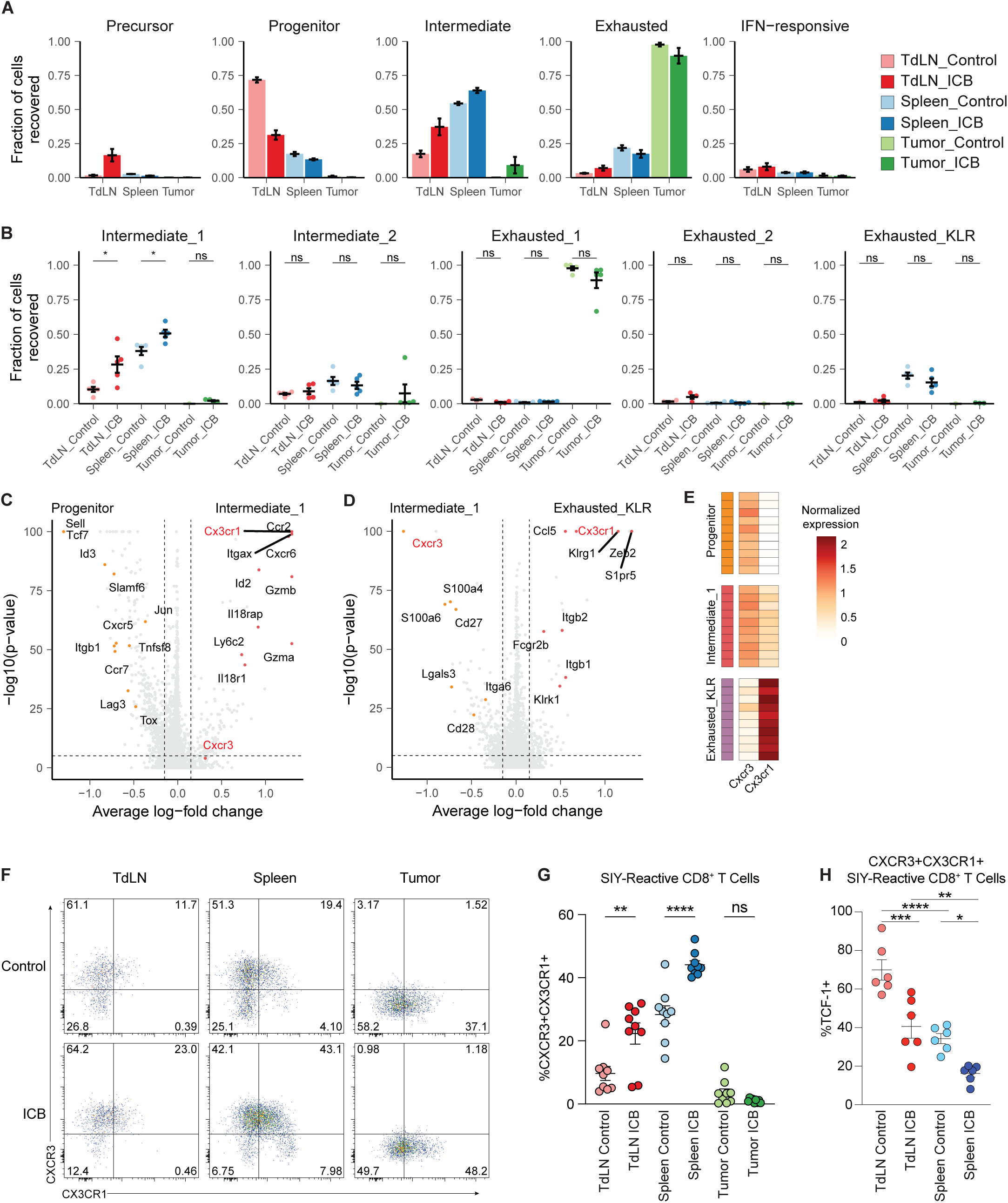
Distribution of transcriptional states between TdLN, spleen, and tumor. A) Frequency of exhausted states in each tissue. B) Frequency of transcriptional states associated with exhaustion in control and ICB-treated mice. P-values are calculated using a two-sided Wilcoxon rank-sum test and are adjusted using Bonferroni correction. C) Volcano plot of transcripts differentially expressed between intermediate_1 and progenitor cells. P-values are calculated using a two-sided Wilcoxon rank-sum test and are adjusted using Bonferroni correction. D) Volcano plot of transcripts differentially expressed between intermrediate_1 and exhausted_KLR cells. P-values are calculated using a two-sided Wilcoxon rank-sum test and are adjusted using Bonferroni correction. E) Heat map of *Cxcr3* and *Cx3cr1* normalized expression by progenitor, intermediate_1, and exhausted_KLR cells. Each cell is the average expression in one mouse. F) Representative flow cytometry plots of SIY reactive T cell expression of CXCR3 and CX3CR1 in TdLN, spleen, and tumor. G) Quantification of CXCR3+CX3CR1+ SIY-reactive CD8+ T cells in TdLN, spleen, and tumor. H) TCF-1 expression of CXCR3+CX3CR1+ SIY- reactive CD8+ T cells in the TdLN and spleen. For G) and H) p-values are calculated using one-way ANOVA.

We also examined the impact of ICB treatment on the frequency of transcriptional states across these three tissues (Figure 3B). We observed that ICB resulted in the expansion of intermediate_1 T cells, but not intermediate_2 T cells, in both the spleen and TdLN. Regardless of ICB treatment, tumors remained nearly entirely comprised of cells with an exhausted_1 phenotype, while exhausted_KLR T cells were found predominantly in the spleen and were largely absent from any other anatomic locations.

To validate the differential expansion of distinct T cell differentiation states in secondary lymphoid organs we established a gating strategy for flow cytometry to distinguish between progenitor, intermediate_1, and exhausted_KLR cells, which comprised the majority of SIY-reactive cells in the TdLN and spleen. We determined which transcripts were differentially expressed among progenitor, intermediate_1, and exhausted_KLR populations (Figure 3C, 3D). We found that progenitor and intermediate_1 T cells expressed comparable levels of *Cxcr3* but that *Cxcr3* expression was absent from exhausted_KLR cells. In contrast, *Cx3cr1* was absent from progenitor T cells, moderately expressed by intermediate_1 T cells, and strongly expressed by exhausted_KLR T cells (Figure 3E). Using flow cytometry, we confirmed the presence of a CXCR3^+^CX3CR1^+^ population, consistent with an intermediate_1 phenotype, as well as a CXCR3^-^ CX3CR1^+^ population, consistent with an exhausted_KLR phenotype, in the spleen of tumor-bearing mice. (Figure 3F). We next assessed the frequency of CXCR3^+^CX3CR1^+^ intermediate_1 cells and CXCR3-CX3CR1^+^ exhausted_KLR T cells. Consistent with scRNA-seq results, we observed an increase in the frequency of CXCR3^+^CX3CR1^+^ intermediate_1 SIY-reactive T cells in both the TdLNs and spleen of mice undergoing treatment with ICB (Figure 3G). Interestingly, the frequency of CXCR3^+^CX3CR1^+^ intermediate_1 SIY-reactive T cells was largest in the spleen. Multiple reports have demonstrated the importance of TCF-1 expression for the ability of CD8^+^ T cells to respond to ICB (1, 20, 23–25, 28); therefore, we also examined the expression of TCF-1 in the CXCR3^+^CX3CR1^+^ intermediate_1 T cell population. We found that a subset of CXCR3^+^CX3CR1^+^ T cells in both the TdLN and spleen expressed TCF-1 (Figure 3H), and that the frequency of TCF-1^+^ T cells decreased in response to ICB, consistent with previous reports (20, 23, 24). Together, these results indicate that the frequency of distinct transcriptional states associated with the progression of T cell exhaustion varies between the tumor, TdLN, and spleen and confirm that treatment with ICB selectively expands T cells of an intermediate_1 phenotype in both the spleen and TdLN.

### CXCR3**^+^**CX3CR1**^+^** intermediate exhausted T cells are derived from progenitor cells and give rise to clonally distinct exhausted_1 or exhausted_KLR phenotypes

To gain insights into the relationships among the expanded T cell clusters, we next examined the TCR repertoire of SIY-reactive cells (Figure 4A). In untreated mice, we observed the largest levels of clonal expansion in the spleen, while in mice treated with ICB, the magnitudes of clonal expansion observed were similar between the tumor, TdLN, and spleen (Figure 4B). Consistent with this observation, we found the diversity of the TCR repertoire decreased in the TdLNs and spleens of mice treated with ICB, and an apparent trend towards reduced diversity in the tumor, indicating that ICB promotes strong expansions of a subset of tumor-reactive clonotypes (Figure 4C). We also observed larger clonal richness (i.e. total number of SIY-reactive clonotypes) in mice treated with ICB in all three tissues surveyed, indicating that ICB treatment simultaneously enables maintenance of a larger repertoire of tumor-reactive T cells (Figure 4D).

**Figure 4:**
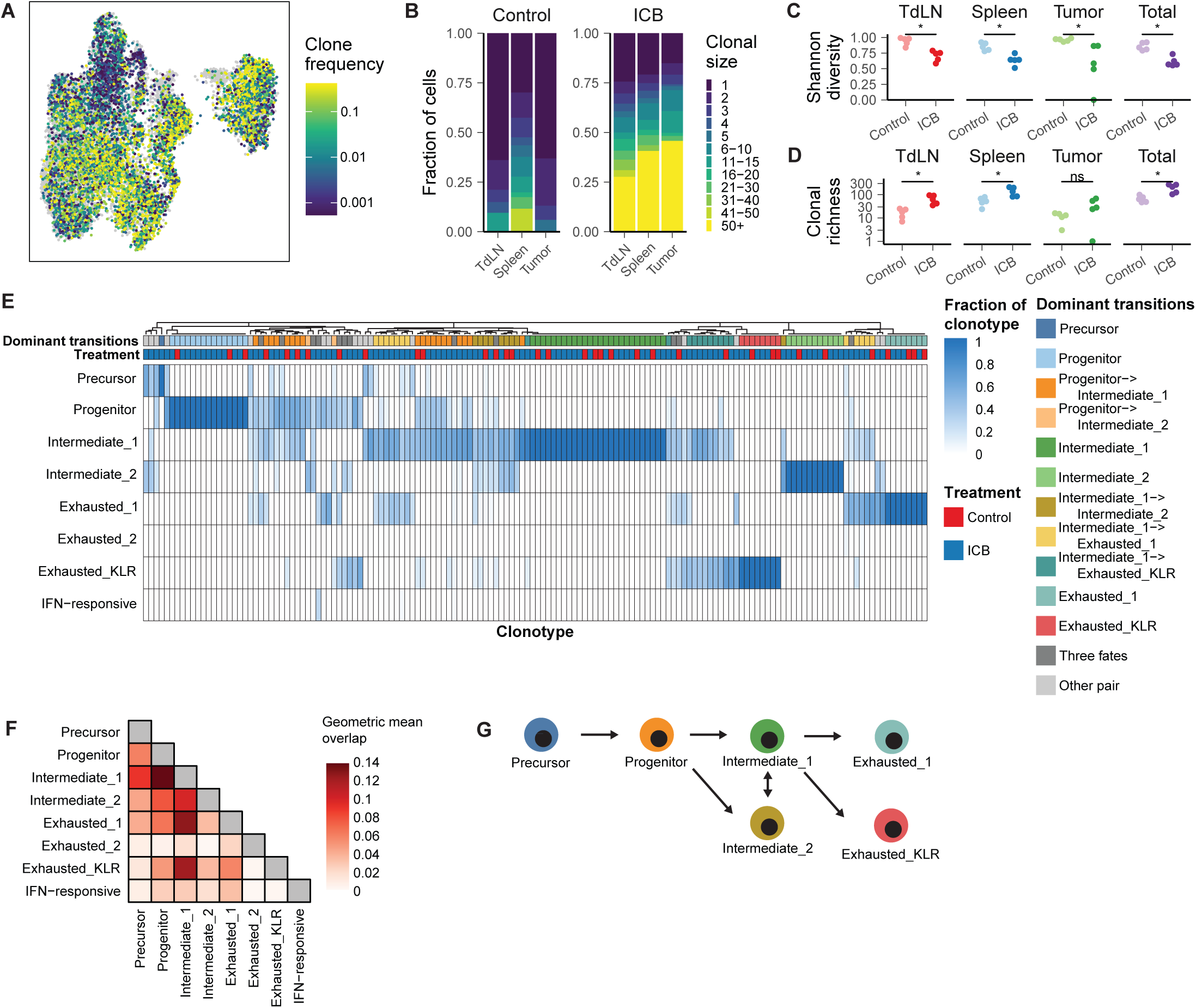
Clonal dynamics of SIY-reactive CD8+ T cells. A) UMAP of SIY-reactive CD8+ T cells colored by clonal size. B) Stacked bar plots of clonal sizes among TdLN, spleen, and tumor, of control and ICB-treated mice. Clonal sizes are computed separately in each tissue. C) Shannon diversity of TCR repertoire. D) Clonal richness of TCR repertoire. E) Heat map of phenotypes present within individual TCR clonotypes. A random sample of the top 150 most expanded clones is shown. Phenotypes present only once within a single clonotype are not shown. F) Transition matrix of transcriptional states. Boxes are shaded according to the geometric mean of normalized clonal frequencies between pairs of transcriptional states. G) Proposed model of T cell differentiation informed by clonal trajectories.

We then sought to define a hierarchy underlying the path of differentiation of tumor-reactive clonotypes. We considered a clonotype “representative” of a phenotype if at least two T cells from that clonotype were found in any given phenotype. Based on this definition, we constructed a heat map of the representative phenotypes present within individual TCR clonotypes (Figure 4E). We then computed a transition matrix using the geometric mean overlap of clonotypes between each pair of phenotypes (Figure 4F). We found that the repertoire of progenitor clonotypes shared substantial overlap with the repertoire of precursor T clonotypes, consistent with precursor T cells undergoing differentiation to progenitor cells. Likewise, intermediate_1 clonotypes exhibited the greatest clonal overlap with progenitor clonotypes, suggesting that intermediate_1 T cells differentiate from progenitor T cells. Intermediate_2 clonotypes demonstrated similar levels of clonal overlap to progenitor clonotypes and intermediate_1 clonotypes, suggesting that progenitor-exhausted T cell clonotypes give rise to both intermediate_1 and intermediate_2 T cells. Both the exhausted_1 and exhausted_KLR phenotypes demonstrated the greatest clonal overlap to intermediate_1 clonotypes, suggesting that the majority of exhausted_1 and exhausted_KLR T cells are generated differentiation of intermediate_1 T cells. In addition, the level of clonal overlap between exhausted_1 and exhausted_KLR phenotypes was minimal, suggesting that they represent distinct, non-interconvertible T cell states. Consistent with this analysis, upset plots depicting the patterns of phenotypic variation within clonotypes demonstrated that the most common pairs of phenotypes identified within a single clonotype were (progenitor, intermediate_1), (intermediate_1, exhausted_1), (progenitor, intermediate_2), (intermediate_1, exhausted_KLR), and (intermediate_1, intermediate_2) (Supplemental Figure 3A). Based on these results, we developed a model in which precursor T cells differentiate into progenitor T cells that in turn give rise to intermediate_1 and intermediate_2 T cells. Intermediate_1 T cells can bifurcate to either exhausted_1 T cells or exhausted_KLR T cells, while intermediate_2 cells appeared to undergo no further detectable differentiation. This model demonstrated high levels of concordance with recent studies of chronic viral infection of mice (33, 34), which included an intermediate phenotype preceding divergent exhausted phenotypes. These observations together suggest that this hierarchy of differentiation represents a fundamental aspect of CD8^+^ T cell biology conserved across multiple antigen reactivities and between chronic viral infection and cancer.

### Differentiation is associated with tissue-site trafficking

We next assessed the relationship between the differentiation of individual clonotypes and their tissue location at the time of analysis. First, we computed the frequency of individual clonotypes in each tissue and found that the frequencies of individual clonotypes among the tumor, TdLN, and spleen were highly correlated, indicating that these three tissues are populated by a common pool of SIY-reactive, tumor-specific clonotypes (Figure 5A). We then identified clonotypes undergoing the phenotypic transitions proposed above. Using the constructed heatmap (Figure 4E), we found the two most common phenotypes encountered within each clonotype, allowing us to classify clonotypes as either belonging to only a single phenotype or transitioning from one phenotype to another (Figure 4E). Clonotypes in which we could not unambiguously assign two dominant phenotypes were excluded from this analysis.

**Figure 5.**
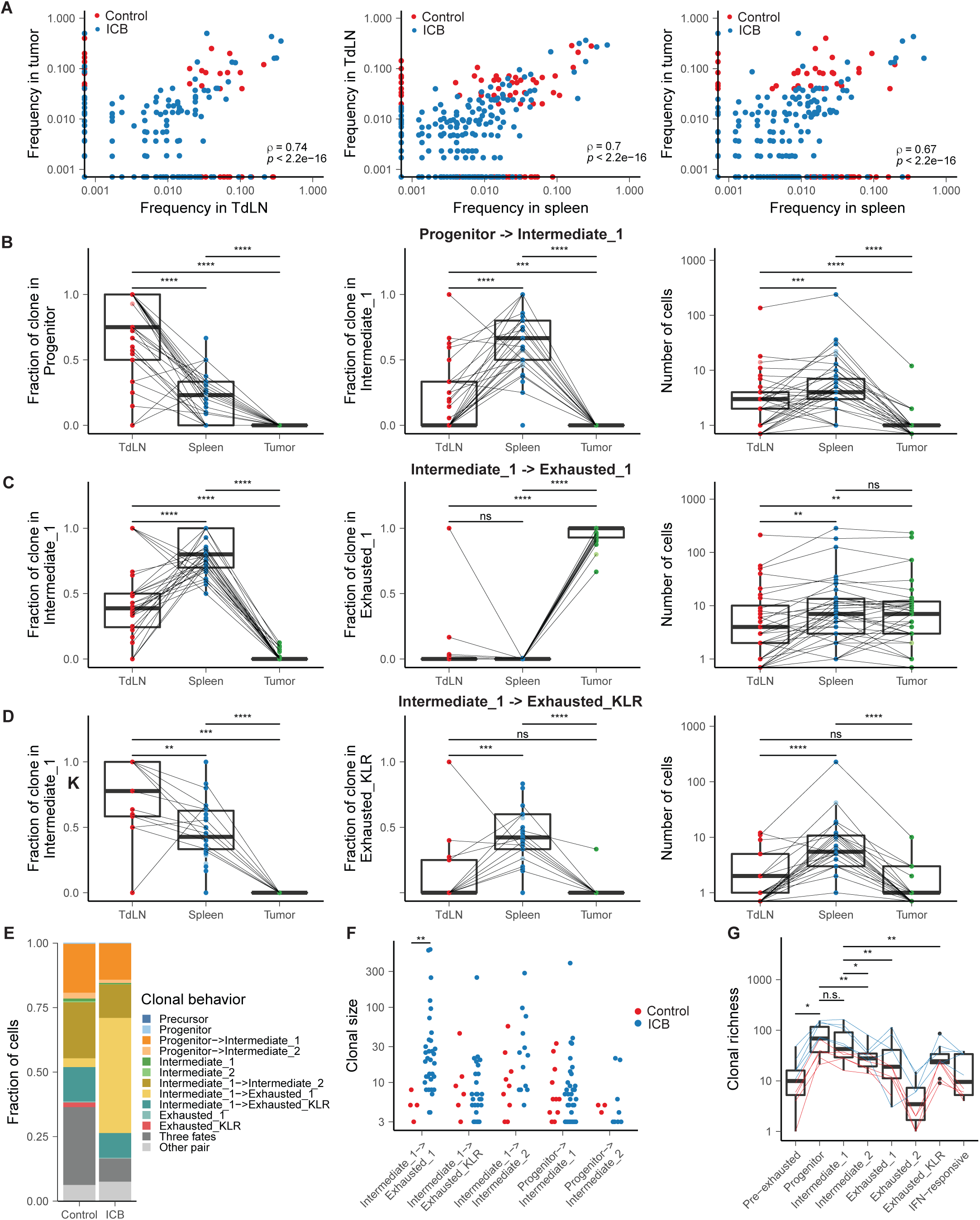
Clonal differentiation is accompanied by change in anatomic site. A) Correlation between clonotype frequencies in the tumor, TdLN and spleen. B) Tissue-site distribution of progenitor and intermediate_1 phenotypes and clonal sizes in progenitor->intermrediate_1 clones. P-values are calculated with a two-sided Wilcoxon rank-sum test and are adjusted with Bonferroni correction. C) Tissue-site distribution of intermediate_1 and exhasuted_1 phenotypes and clonal sizes in intermediate_1 -> exhausted_1 clones. P-values are calculated with a two-sided Wilcoxon rank-sum test and are adjusted with Bonferroni correction. D) Tissue-site distribution of intermediate_1 and exhausted_KLR phenotypes and clonal sizes in intermediate_1 -> exhausted_KLR clones. P-values are calculated with a two-sided Wilcoxon rank-sum test and are adjusted with Bonferroni correction. E) Stacked bar chart of clonal behaviors assigned to single cells in control and ICB-treated mice. Only clonotypes consisting of more than two cells are shown. F) Absolute clonal sizes of transitioning clonotypes in control and ICB-treated mice. P-values are calculated with a two-sided Wilcoxon rank-sum test and are adjusted with Bonferroni correction. G) Clonal richness of transcriptional phenotypes in control and ICB-treated mice. P-values are calculated with a paired, two-sided Wilcoxon rank-sum test and are adjusted with Bonferroni correction.

We then compared the clonal sizes and phenotypes present at each tissue site within transitioning clonotypes. We focused this analysis on clonotypes undergoing transitions between progenitor, intermediate_1, intermediate_2, exhausted_1, and exhausted_KLR phenotypes, which accounted for the majority of clonal transitions observed (Figure 4E). We found that clonotypes differentiating from the progenitor state to intermediate_1 were strongly polarized towards the progenitor phenotype in the lymph node but were polarized towards the intermediate_1 phenotype in the spleen, indicating that these transition between two transcriptional states is strongly related to the migration of the T cells from the TdLN to the spleen (Figure 5B, Supplemental Figure 3B). Likewise, the transition between intermediate_1 and exhausted_1 phenotypes was strongly associated with trafficking from the spleen to the tumor (Figure 5C). In contrast, clonotypes undergoing differentiation from intermediate_1 to exhausted_KLR or from intermediate_1 to intermediate_2 were largely absent from the TdLN and the tumor but present in the spleen, indicating that these transitions primarily occur within the spleen and are not accompanied by transit to a different tissue site (Figure 5D, Supplemental Figure 3C). Overall, these results demonstrate that clonally related T cells possess distinct phenotypes in different tissues and that select phenotypic transitions, including progenitor è intermediate_1, progenitor è intermediate_2 and intermediate_1 è exhausted_1, are strongly associated with trafficking from one tissue to another, while others, such as intermediate_1 è exhausted_KLR and intermediate_1 è intermediate_2 are not accompanied with a change in tissue site and instead take place primarily among cells resident in the spleen.

### The intermediate_1 → exhausted_1 transition limits clonal differentiation and is modulated by checkpoint blockade

To determine how ICB may modulate the phenotypic transitions experienced by SIY-reactive clonotypes, we examined the distribution of clonal behaviors present in control and ICB-treated mice. Although the frequency of clonotypes classified as transitioning between intermediate_1 and exhausted_1 was similar, the fraction of T cells belonging to an intermediate_1 → exhausted_1 transitioning clonotype was substantially greater in mice treated with ICB (Figure 5E, Supplemental Figure 3D). We then calculated the absolute clonal sizes of individual clonotypes in control and ICB-treated mice undergoing each of the five clonal transitions analyzed above (Figure 5E, Supplemental Figure 3D). We also found that the clonal sizes of intermediate_1 → exhausted_1 clonotypes, but no other set of transitioning clonotypes, were significantly larger in mice treated with ICB, suggesting that ICB preferentially expands intermediate_1 clonotypes that are primed to undergo an intermediate_1 to exhausted_1 transition (Figure 5F).

We also computed the clonal richness of each phenotype encountered on our hypothesized differentiation trajectory (Figure 5G). We found that clonal richness peaked at the progenitor and intermediate_1 phenotypes, suggesting that precursor cells, the apparent direct precursors of progenitor cells, are largely depleted by day 14 following tumor inoculation. In addition, this result demonstrates that there is an accumulation of clonotypes at the progenitor and intermediate_1 phenotypes. This result suggests that while the transition from progenitor to intermediate_1 phenotype is efficient – that is, it occurs with high probability--the transition from intermediate_1 to an exhausted_1 or exhausted phenotype is inefficient, and that the intermediate_1 phenotype is a rate-limiting step encountered during clonal differentiation. Thus, on the basis of our data, we concluded that one of the major results of ICB treatment is the expansion of splenic intermediate_1 clonotypes that are pre-disposed to undergo the intermediate_1 → exhausted_1 transition.

### Splenic Intermediate_1 T cells drive the expansion of tumor-reactive CD8^+^ T cells during ICB

To confirm whether the ability of tumor-reactive CD8^+^ T cells to expand in response to ICB was different between T cells from TdLN and spleen, we used an adoptive transfer approach of TCR transgenic 2C T cells (Figure 6A). In this system, naïve 2C T cells become rapidly activated in response to KP.SIY tumors, and expanded 2C T cells appear in both the TdLNs and spleens of mice within 72 hours after adoptive transfer. We first investigated the phenotypes of transferred 2C T cells in TdLNs and spleens. Consistent with our previous data, significantly more 2C T cells in the spleen showed an intermediate_1 CXCR3^+^CX3CR1^+^ phenotype compared to 2C cells in the TdLN. This observation supports that differentiation from progenitor to Intermediate_1 is associated with splenic trafficking (Figure 6B, 5C).

**Figure 6:**
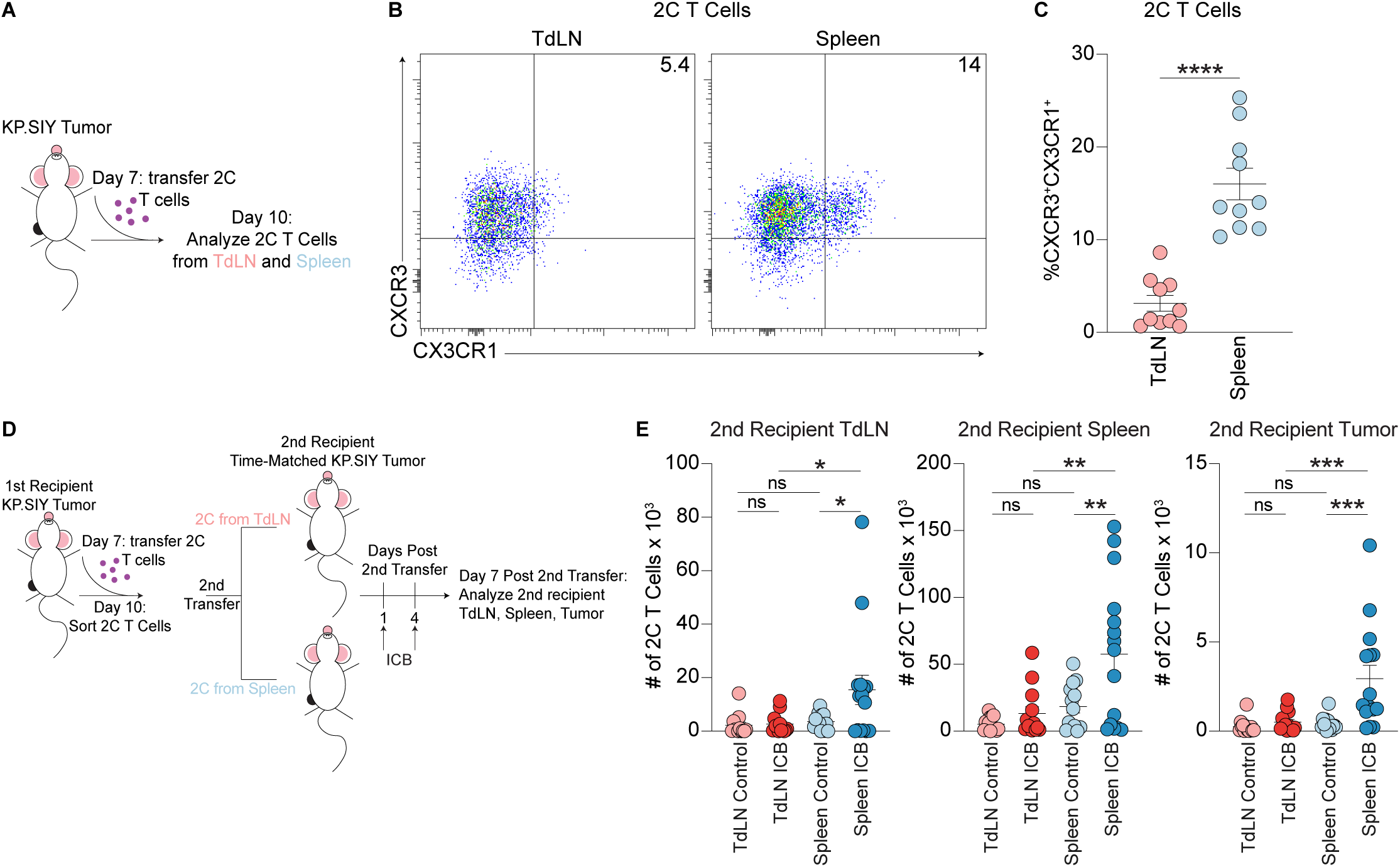
Splenic tumor-reactive CD8+ T cells drive the response to ICB. A) Adoptive transfer experimental design. B) Phenotype of 2C T cells 3 days after transfer to KP.SIY tumor-bearing mice. C) Quantification of the phenotype of 2C T cells 3 days after transfer to KP.SIY tumor bearing mice, n = 10. P-value calculated with a Mann-Whitney U test. D) Experimental design of transfer into secondary recipients. E) Accumulation of transferred 2C T cells in secondary recipients. TdLN Control n = 13, TdLN ICB n=13, Spleen control n=14, Spleen ICB n=15. P-values calculated with one-way ANOVA.

We next determined whether trafficking to the spleen was associated with increased responsiveness to ICB. We sorted 2C T cells from TdLNs and spleens of primary recipient mice and transferred the sorted 2C T cells to secondary recipients with time-matched KP.SIY tumors (Figure 6D). Strikingly, only 2C T cells sorted from spleens of mice displayed expansion in response to ICB (Figure 6E). This expansion occurred not only in spleens of secondary recipients, but also the TdLNs and tumors (Figure 6E). Our interpretation of these results is that intermediate_1 T cells in the spleen are primary responders to ICB, and that intermediate_1 T cells are able to recirculate to both TdLNs and tumors. Recirculation of expanding intermediate_1 T cells from the spleen to TdLNs therefore appears to explain the increased frequency of tumor-reactive T cells observed in the TdLNs following ICB.

### Antigen Density Regulates Intermediate_1 Differentiation and Trafficking

We sought to further understand the regulators of tumor-reactive CD8^+^ T cell differentiation in the spleen. Chronic antigen stimulation is known to drive T cells into terminal exhaustion states (12, 13). Consistent with this notion, SIY-reactive CD8^+^ T cells infiltrating the tumor were almost completely comprised a terminally exhausted phenotype. Progenitor exhausted CD8^+^ T cells are also induced by high antigen levels in the lymph node (18), but how antigen affects differentiation of T cells into the intermediate_1 phenotype is unknown. In the tumor, antigen is abundant via direct presentation from tumor cells or cross presentation from APCs. Outside of the tumor environment, antigen presentation to T cells is reliant on APCs. To examine the extent of antigen trafficking to the spleen, we inoculated mice with KP cells that expressed the pH-stable fluorophore zsGreen (KP.zsGreen), and examined the levels of zsGreen in CD45^+^ cells in TdLNs and spleens. TdLNs had a significantly higher frequency of zsGreen^+^CD45^+^ cells compared to spleens, indicating that the spleen was a relatively antigen low anatomic site (Figure 7A). Using the 2C adoptive transfer approach, we then determined whether antigen was required to drive tumor-reactive CD8^+^ T cell expansion in the spleen. We primed 2C T cells in vivo in primary recipient KP.SIY tumor-bearing mice, and then transferred 2C T cells isolated from the spleens of primary recipients to secondary hosts bearing either KP.SIY or KP tumors. Secondary hosts were treated with ICB (Figure 7B). 2C T cells failed to accumulate in secondary hosts bearing KP tumors, indicating that antigen was required for 2C T cell expansion in response to ICB, even at low levels (Figure 7C).

**Figure 7:**
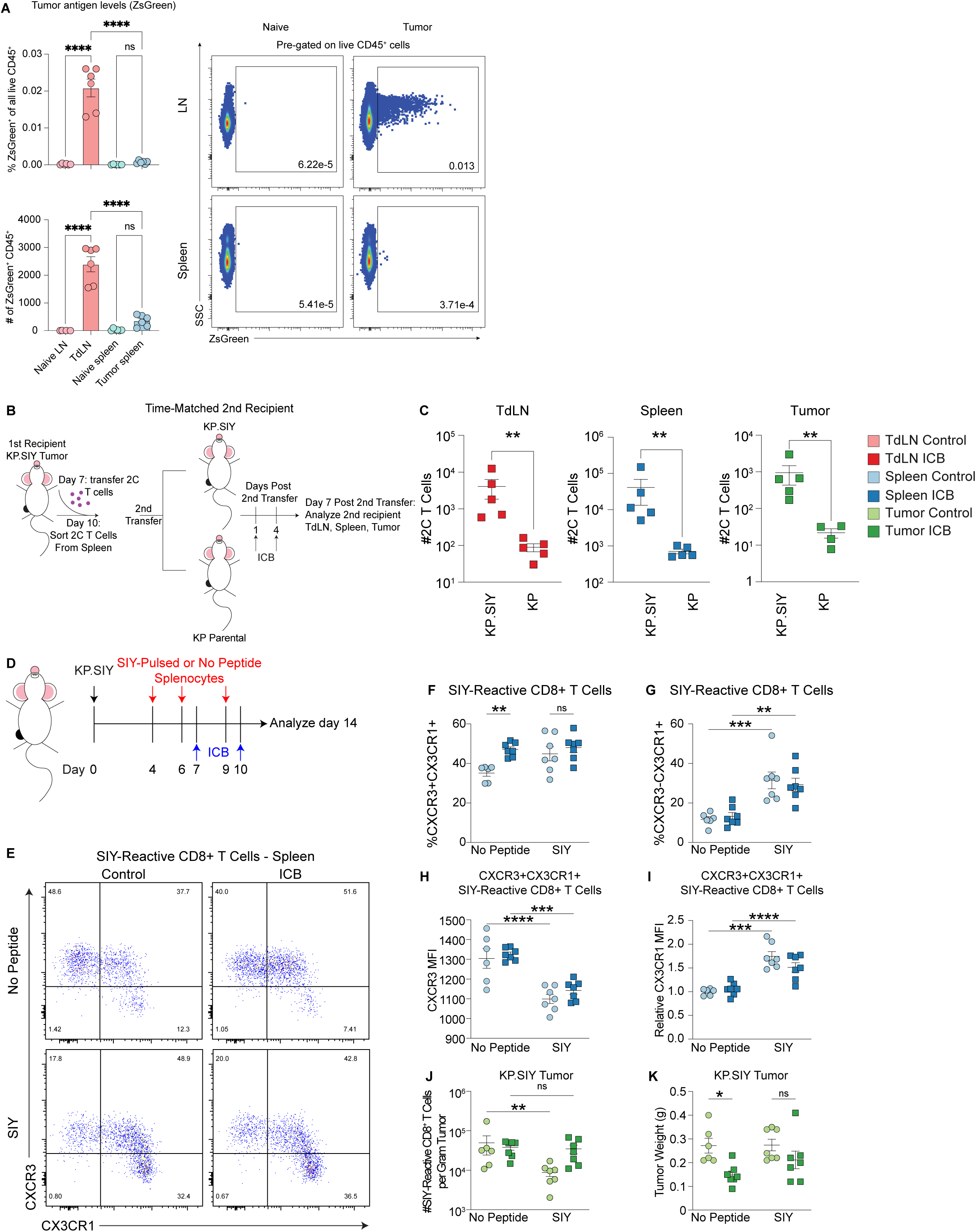
Antigen levels impact SIY-reactive T cell differentiation in the spleen and trafficking to the tumor. A) Tumor-derived Zsgreen levels in TdLNs and spleens, n=6. P-values calculated with one-way ANOVA. B) Experimental scheme of 2C adoptive transfer. C) Numbers of recovered 2C T cells in secondary recipients with either KP.SIY or KP tumors, n=5 (or 4 for KP tumor). P-values calculated with Mann-Whitney U test. D) Experimental Scheme of SIY-pulsed splenocyte transfer. D) Example flow cytometry plots of SIY-reactive T cell expression of CXCR3 and CX3CR1 in the spleen F) Percentage of splenic SIY-reactive CD8+ T cells that are CXCR3+CX3CR1+. G) Percentage of splenic SIY-reactive CD8+ T cells that are CXCR3-CX3CR1+. H) CXCR3 MFI of splenic SIY-reactive CXCR3+CX3CR1+ CD8+ T cells. I) Relative CX3CR1 MFI of splenic SIY- reactive CXCR3+CX3CR1+ CD8+ T cell population. J) Number of SIY-reactive CD8+ T cells per gram of KP.SIY tumor. K) KP.SIY tumor weights. For F-K, n=6, p-values calculated with 2-way ANOVA.

We then aimed to determine whether increased delivery of antigen to the spleen would modulate T cell differentiation. We isolated splenocytes from naïve mice, pulsed them with SIY, and transferred them to KP.SIY tumor-bearing mice, which were subsequently treated with ICB (Figure 7D). Examining mice on day 14 of tumor growth, we found that delivery of SIY-pulsed splenocytes had no effect on the proportion of Intermedate_1 (CXCR3^+^CX3CR1^+^) T cells in the spleen (Figure 7E and 7F), but resulted in an increased frequency of exhausted_KLR T cells (CXCR3^-^CX3CR1^+^) (Figure 7E and 7G). Additionally, T cells with the CXCR3^+^CX3CR1^+^ Intermediate_1 phenotype expressed lower levels of CXCR3 and higher levels of CX3CR1 in mice that received SIY-pulsed splenocytes (Figure 7H and 7I). Intermediate_1 T cells in the spleen therefore appeared to undergo differentiation to the exhausted_KLR phenotype in response to encounter of antigen at high density. This shift in T cell differentiation in the spleen to the exhausted_KLR phenotype in response to SIY-pulsed splenocytes also associated with a striking reduction of tumor-infiltrating SIY-reactive CD8^+^ T cells and failed tumor control following ICB (Figure 7I, 7J). These results suggest that low but detectable levels of antigen in the spleen are critical for the accumulation and retention of T cells in the intermediate_1 state, which can traffic to the tumor and further differentiate into exhausted_1 T cells. In contrast, high antigen levels drives differentiation of intermediate_1 cells into the exhausted_KLR phenotype. Consistent with our previous observation that exhausted_KLR T cells resided primarily in the spleen (Figure 3B), exhausted_KLR differentiation in response to SIY-pulsed splenocytes hampered tumor trafficking, resulting in decreased numbers of antigen-specific T cells in the tumor microenvironment (Figure 7J). Successful ICB therefore appears to rely on low antigen levels in the spleen to allow for the accumulation of Intermediate_1 T cells that can traffic to the tumor.

### Exhausted_KLR T cells in human patients exhibit reduced migration to tumors relative to other tumor-homing clonotypes

To assess the extent to which the transcriptional phenotypes and clonotypic trends observed among SIY-reactive T cells in our mouse model are present in human patients, we conducted a reanalysis of a pan-cancer atlas of tumor-infiltrating lymphocytes recovered from 316 cancer patients (37). We focused this analysis on a subset of the larger atlas, comprising four types of solid tumors (colorectal carcinoma, hepatocellular carcinoma, non-small cell lung cancer, and cholangiocarcinoma) for which single-cell RNA and TCR sequencing data from matched tumor tissue and peripheral blood were available. On the basis of transcriptional expression, the authors annotated the majority of CD8^+^ T cells as either: naïve (Tn), memory (Tm), resident memory (Trm), effector memory (Tem), exhausted (Tex), terminally differentiated memory or effector cells (Temra), NK-like (Tnk), or ISG-positive (ISG).

We first examined the concordance between SIY-reactive transcriptional phenotypes identified in our mouse model and the transcriptional phenotypes present among TIL from human patients. We found strong agreement between the signatures identified in our mouse model and the phenotypes annotated by Zheng et al: specifically, Tn expressed strong levels of our progenitor signature, Tm, Trm, and Tem upregulated our intermediate_1 signature, with a subset of Trm also upregulating our intermediate_2 signature, Tex upregulated our exhausted_1 and exhausted_2 signatures, and Temra upregulated our exhausted_KLR signature (Figure 8A-C, Supplemental Figure 4). Thus, the transcriptional states present in our mouse model can differentiate phenotypes of T cells present in the tumor and peripheral blood of human patients, and Temra are the most analogous coutnerpart of the exhausted_KLR cells that we have identified.

**Figure 8.**
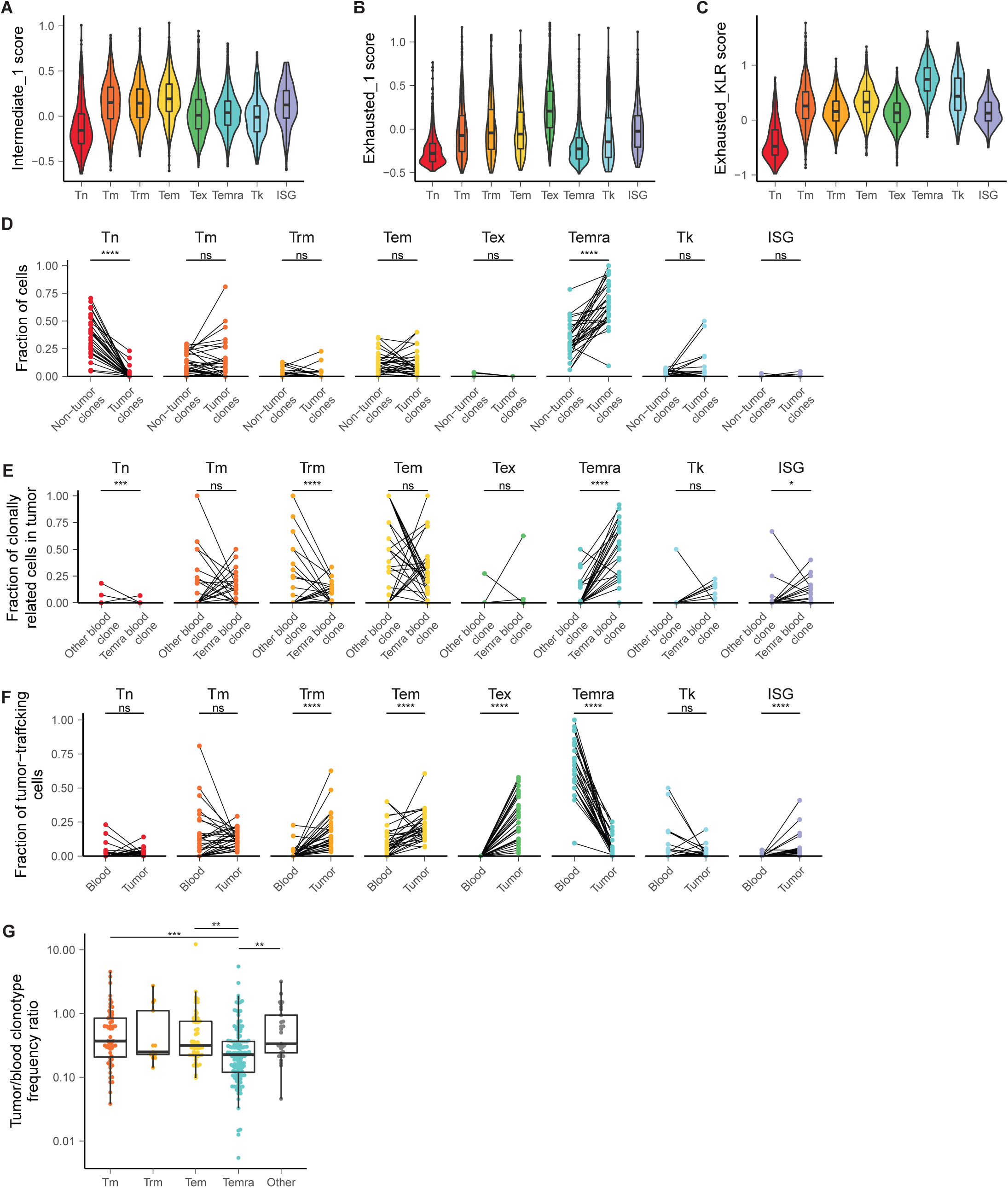
Exhausted_KLR cells in human patients exhibit decreased migration from peripheral blood to tumor. A-C) Expression of signatures for intermediate_1, exhausted_1, and exhausted_KLR phenotypes on clusters defined by Zheng et al. D) Frequency of phenotypes present in peripheral blood among tumor-trafficking and non-tumor-trafficking clonotypes. P-values are calculated with a two-sided Wilcoxon rank-sum test and are adjusted with Bonferroni correction. E) Frequency of phenotypes present among tumor-infiltrating cells related to Temra clonotypes from peripheral blood and all other tumor-infiltrating clonotypes. P-values are calculated with a two-sided Wilcoxon rank-sum test and are adjusted with Bonferroni correction. F) Frequency of phenotypes present among tumor-trafficking CD8 T cells in blood and tumor. P-values are calculated with a two-sided Wilcoxon rank-sum test and are adjusted with Bonferroni correction. G) Ratio of clonal frequency in tissue to clonal frequency iamong Tm, Trm, Tem, Temra, and all other clonotypes. P-values are calculated by a Kruskal-Wallis rank-sum test followed by Dunn’s post-test.

Next, we analyzed T cells in the peripheral blood that were clonally related to tumor infiltrating clonotypes. We found that Temra cells in the peripheral blood were the only population increased in frequency among tumor-trafficking clonotypes relative to non-tumor-trafficking clonotypes, suggesting that this population in peripheral blood is enriched for tumor-specific CD8^+^ T cells (Figure 8D). We also found that the tumor-infiltrating clonal relatives of Temra cells from the peripheral blood were also enriched for a Temra phenotype, relative to other tumor-infiltrating clonotypes found in peripheral blood, demonstrating that this phenotype is conserved upon entry into the tumor (Figure 8E).

To determine whether trafficking of T cells into the tumor was associated with any specific T cell state we assessed the faction of T cells within each state between tumor and peripheral tissues. Strikingly, we found that among tumor-trafficking clonotypes, the Temra phenotype was substantially decreased in frequency in the tumor relative to peripheral blood, suggesting poor tumor infiltration (Figure 8F). In contrast, Trm, Tem, and ISG cells exhibited an increase in frequency in the tumor, suggesting that tumor-specific Trm, Tem, and ISG cells present in the peripheral blood undergo efficient recruitment to the tumor. To assess whether these trends in tumor trafficking potential were apparent at the level of individual clonotypes, we computed the ratio between the frequency of each clonotype in the tumor and the blood. We found that relative to other clonotypes in the peripheral blood, Temra clonotypes were substantially more expanded in peripheral blood compared to the tumor, indicating that individual Temra clonotypes exhibit a reduced ability to enter the tumor, relative to other phenotypes in the peripheral blood (Figure 8G). Overall, these data support our model in which intermediate_1-like T cells (Tm, Trm, and Tem) undergo efficient entry into the tumor from peripheral tissues, while exhausted_KLR-like clonotypes (Temra) exhibit a decreased potential to migrate to the tumor, especially relative to their level of clonal expansion in peripheral blood.

## Discussion

Here, we analyzed tumor-reactive CD8^+^ T cells from the tumor, TdLN, and spleen of ICB-treated and untreated mice with scRNA and TCR sequencing. Our findings corroborate previous studies that demonstrated the existence of distinct transcriptional phenotypes associated with CD8^+^ T cell exhaustion, including multiple intermediate and exhausted CD8^+^ T cell differentiation states (33, 34). We extend prior knowledge of these differentiation states by demonstrating that these transcriptional phenotypes appear largely restricted to individual anatomic sites in tumor-bearing mice. Additionally, we found that the splenic intermediate_1 population gives rise to two terminally differentiated populations of CD8^+^ T cells. One of these populations, exhausted_1, comprises the majority of tumor-infiltrating clones, including the clones that expand in response to ICB, and becomes canonically exhausted in the tumor microenvironment. The splenic intermediate_1 population also gives rise to a largely clonally distinct exhausted_KLR population of CD8^+^ T cells that remains in the spleen and traffics to the tumor with low efficiency. We also found that increased antigen density in the spleen increased the differentiation of intermediate_1 to the exhausted_KLR state, suggesting that alternative states of exhaustion emerge depending on the anatomic site of chronic antigen encounter. Increased differentiation to an exhausted_KLR state reduced the number of tumor-infiltrating T cells and the efficacy of checkpoint blockade, suggesting that maintaining a pool of intermediate_1 T cells in the spleen, capable of expanding in response to ICB and subsequently trafficking to the tumor, is critical for immunotherapy efficacy. Lastly, we found that human T cells expressing gene signatures concordant with our exhausted_KLR population appear to expand in response to tumor antigen but traffic to the tumor with low efficiency. Together, our results demonstrate that microenvironmental factors present in individual anatomic sites play a role in guiding exhausted T cell differentiation and are critical in determining the outcome of ICB.

The spleen is frequently analyzed in models of chronic viral infection, including LCMV, but in these models, the spleen is also a primary site of infection. In contrast, the spleen is not often studied in cancer (20, 23–25, 27, 28). We found that splenic intermediate_1 T cells exhibit the greatest expansion in response to ICB; consistent with this result, Giles et al. also recently proposed that intermediate exhausted cells are most expanded by anti-PD-L1 treatment in murine LCMV infection (33). We found that a subset of intermediate_1 T cells express TCF-1, consistent with previous reports identifying TCF-1^+^ CD8^+^ T cells as the mediators of T cell self-renewal and expansion in response to ICB (1, 20, 23–25, 28). We propose that the accumulation of TCF1^+^ progenitor CD8^+^ cells in TdLNs involves the expansion of intermediate_1 T cells in the spleen and their subsequent recirculation from the spleen to the TdLN. In support of this, we find that the transition from the progenitor to the intermediate_1 transcriptional phenotype correlates with entry to the spleen. Additionally, we find that adoptively transferred 2C cells from the spleen, but not TdLN, exhibit an enhanced ability to expand in secondary recipients upon treatment with ICB, and result in increased numbers of 2C cells in the tumor, TdLN, and spleen, demonstrating the ability of splenic CD8^+^ T cells to recirculate to the TdLN.

Intermediate_1 T cells also bear some resemblance to a population of “transitory” proliferating cells reported to express CX3CR1 (23, 38). These studies have suggested that these transitory T cell derived directly from CXCR5^+^ progenitor T cells; our data suggests the majority of ICB- triggered expansion originated from intermediate_1 → exhausted_1 cells. A reported population of CXCR5^+^ CD8^+^ progenitor exhausted T cells that proliferated after blockade of PD-L1 also expressed CXCR3 has strong similarities to similar to intermediate_1 T cells (24). It may be that in cancer, the intermediate_1 population combines aspects of progenitor and transitory T cells found in chronic LCMV infection. Intermediate_1 T cells therefore likely have a greater role in the response to ICB than previously appreciated, especially in cancer.

Similar to previous studies (33, 34), we identified that the intermediate_1 population served as a hub for differentiation into multiple phenotypic states, including intermediate_2, exhausted_1, and exhausted_KLR. Of these phenotypes, only exhausted_1 exhibited efficient trafficking of the tumor, suggesting that state decisions made at this bottleneck substantially impact the effectiveness of an anti-tumor T cell response. These three T cell states also exhibited minimal clonal overlap, suggesting that they represent divergent fates, and raising the possibility that TCR signaling characteristics may influence these fate decisions. Indeed, prior studies have demonstrated that higher TCR affinity promotes differentiation into a terminally exhausted phenotype, rather than a phenotype resembling exhausted_KLR (34). Future studies could seek to understand factors that regulate the fate commitment of intermediate_1 T cells. Understanding how to effectively target and manipulate the intermediate_1 → exhausted_1 fate decision may offer new strategies to increase the tumor-infiltrating abilities of T cells and enhance the efficacy of ICB.

By examining the relationships between anatomic sites and transcriptional phenotypes within individual clonotypes, we observed that some phenotypic transitions, including progenitor → intermediate_1 and intermediate_1 → exhausted_1, are strongly associated with transit between different tissues. We are unable to conclude whether entry into different tissues is a key factor promoting differentiation, or whether differentiation itself promotes transit to different tissue sites. One previous study found, however, that full effector differentiation of progenitor T cells requires entry into the tumor, suggesting that T cell state transitions can result from trafficking to new anatomic locations (39).

In contrast to intermediate_1 T cells, our results indicate that mouse and human CX3CR1^hi^ CD8^+^ T cells, including exhausted_KLR and Temra T cells, respectively, do not traffic efficiently to the tumor. This result is consistent with previous findings where CX3CR1^hi^ memory CD8^+^ T cells were preferentially found in blood, while CXCR3^+^CX3CR1^int^ memory T cells, which phenotypically resemble intermediate_1 cells, were resident in non-lymphoid tissues (40). These differential homing patterns may be explained by the expression of CXCR3; intermediate_1 T cells express significantly more CXCR3 than exhausted_KLR cells, and CXCR3 expression is critical for the homing of CD8^+^ T cells into tumors (41). Future studies could focus on the contributions of exhausted_KLR CD8^+^ T cells to anti-tumor immunity. Consistent with our finding of inefficient tumor homing by exhausted_KLR T cells, two previous studies using subcutaneous mouse models of cancer found that CX3CR1^hi^ CD8^+^ T cells contribute little to anti-tumor immunity (42, 43). One could imagine, however, that a population of circulating or spleen-patrolling tumor-reactive CD8^+^ T cells could surveil for disseminated tumor cells and protect against metastasis (44). These populations could also potentially provide a blood-based biomarker for anti-tumor immunity or response to ICB (45).

Previous studies of memory responses following acute infection and chronic infection, including mouse models of MCMV and *T. gondii* infection, have highlighted important roles for CXCR3- expressing “intermediate” CD8^+^ T cell populations (46–48). We find that splenic antigen levels are important in guiding the differentiation of intermediate_1 T cells. Similar to the tumor model observed here*, T. gondii* infection exhibits a pattern of tissue-restricted antigen presence, with high antigen burden in the brain, but low antigen burden in other tissues, including the spleen (46). We propose that these patterns of tissue-restricted antigen presence may encourage profound yet transient antigen encounter in infected or tumor tissue, or their draining lymph nodes, followed by low density antigen encounter in the spleen. This pattern of antigen encounter may represent a key component of the anti-tumor immune response that is not well-modeled by the LCMV model of chronic viral infection, which involves high antigen levels in the spleen as well in many other tissues throughout the organism. Recent studies of CAR-T cells have demonstrated that transient rest from antigen encounter leads to enhanced anti-tumor efficacy and results in the reversal of T cell exhaustion (49–51), suggesting that transient or intermittent antigen encounter is a key factor for sustaining the functionality of an anti-tumor T cell response.

One limitation of our study is that there remains insufficient available data from tumor-reactive CD8^+^ T cells recovered from the TdLNs and spleens of human cancer patients to assess to what extent the transcriptional states and phenotypic transitions that we identified in our mouse models reflect patterns of differentiation experienced by anti-tumor CD8^+^ T cells in human cancer patients.

Together, our data highlight a previously unappreciated role for the spleen in coordinating the differentiation of tumor-reactive CD8^+^ T cells as they respond to ICB. The splenic intermediate_1 phenotype that we identified comprised the majority of tumor-reactive CD8^+^ T cells that expanded upon ICB treatment and gave rise to a majority of tumor-infiltrating clonotypes, suggesting that is a crucial part of the anti-tumor immune response. This study provides mechanistic insight into responses to ICB and will inform future studies of anti-tumor immune responses in cancer.

## Materials and Methods

### Mice

Male and female C57BL/6 mice were obtained from Taconic Biosciences or the Jackson Laboratories. Male and female 2C Rag2^-/-^CD45.1^+^ mice were bred and maintained in house. All mice were housed and bred under specific pathogen free (SPF) conditions at the Koch Institute animal facility. Mice were gender-matched and age-matched to be 6-12 weeks old at the time of experimentation. All experimental animal procedures were approved by the Committee on Animal Care (CAC/IACUC) at MIT.

### Tumor cell lines and tumor outgrowth studies

The parental KP NSCLC cell line was a gift from the Jacks laboratory at MIT. The KP tumor line stably expressing cerulean-SIY and the KP tumor line stably expressing zsgreen were generated previously (Horton et. al. Science Immunology 2021). Expression of fluorescent molecules was periodically assessed using flow cytometry. Tumor cell lines were cultured at 37°C and 5% CO_2_ in DMEM (Gibco) supplemented with 10% FBS (Atlanta Biologicals), 1% penicillin/streptomycin (Gibco), and 1X HEPES (Gibco). Tumor cells were harvested by trypsinization (Gibco) and washed 2 times with 1X PBS (Gibco). Cells were resuspended in PBS, and 2.5×10^5^ tumor cells were injected subcutaneously into the flanks of mice. Subcutaneous tumor area measurements (calculated as *length* x *width*) were collected 2-3 times a week with calipers until the endpoint of the study.

### ICB treatment

Mice were injected intraperitoneally with anti-CTLA-4 (clone UC10-4F10-11, Bio X Cell) and anti-PD-L1 (clone 10F.9G2, Bio X Cell) antibodies on days 7, 10, 13, and 16 post-tumor inoculation for tumor outgrowth studies, or on days 7 and 10 post-tumor inoculation for day 14 analyses. Each mouse received 100 μg of each antibody per treatment.

### Tumor and tissue dissociation

To discriminate vascular T cells from T cells in tissues, tumor-bearing mice were injected retro-orbitally with fluorescently-labeled anti-CD45 antibodies (CD45-IV) 3 minutes prior to euthanasia. Spleens and lymph nodes were dissected from mice and physically dissociated through a 70 µm filter to generate single cell suspensions. Splenocyte suspensions were lysed with ACK lysis buffer (Gibco) for 3 minutes to deplete red blood cells. Subcutaneous tumors were dissected from mice, weighed, and collected in 5 mL DMEM (Gibco) containing the human tumor dissociation kit (Miltenyi) enzymes. Tumors were dissociated using a Mixer HC (USA Scientific) set to 37 degrees Celsius, shaking at 500 RPM for 45 – 60 minutes. Following the digestion, tumors were mashed through a 70 μm filter with a 1 mL syringe plunger to generate a single cell suspension. The dissociated cells were washed 2 times with PBS and were layered over Ficoll (GE). Cells were spun over Ficoll at 450 *g* for 30 minutes with the lowest settings of acceleration and brakes. The layer at the interface of Ficoll and PBS was collected and washed with PBS.

### Flow cytometry and cell sorting

Prior to staining, cells were washed with FACS staining buffer (chilled PBS containing 1% FBS and 2 mM EDTA). Cells were resuspended in 50 μL of the antibody-containing staining buffer, plus eBioscience Fixable Viability Dye eFluor 780 or eFluor 506 to distinguish live and dead cells and with anti-CD16/CD32 (clone 93, BioLegend) to prevent non-specific antibody binding. Cell surface proteins were stained for 20 min on ice with fluorophore-conjugated antibodies at a 1:200 dilution. Cells were then washed twice and resuspended in eBioscience Fixation/Permeabilization buffer and incubated 30 minutes at room temperature. Cells were then washed twice and resuspended in staining buffer with intracellular antibodies. To obtain absolute counts of cells, Precision Count Beads (BioLegend) were added to samples according to manufacturer’s instructions. Flow cytometry sample acquisition was performed on a LSR Fortessa cytometer (BD), and the collected data was analyzed using FlowJo v10.5.3 software (TreeStar). For cell sorting, the surface staining was performed as described above under sterile conditions, and cells were acquired and sorted into complete medium using a FACSAria III sorter (BD). For CD8^+^ T cell analysis, cells were pre-gated on FSA and SSC, Live, CD45^+^, CD45-IV^-^, CD3^+^ or TCRbeta^+^, single cells, CD4^-^, CD8^+^. Antibodies used: CD4 clone RM4-5, CD45 clone 30-F11, CD45.1 clone A20, CD45.2 clone 104, CD8a clone 53-6.7, TCRβ clone H57-597, CD3 clone 17A2, Thy1.2 clone 53-2.1, CXCR3 clone CXCR3-173, CX3CR1 clone SAO11F11, KLRG1 clone 2F1/KLRG1. SIY-tetramer was obtained from the NIH tetramer core, conjugated to PE, and added at 1:500-1:1,000 to the extracellular staining antibody mix. For scRNA-seq experiments with cell hashing, cells were also stained with Totalseq A anti-mouse hashing antibodies (Biolegend) before FACS.

### 2C T cell adoptive transfer

Spleens and inguinal lymph nodes of 2C RAG2^-/-^CD45.1^+^ mice were dissected and made into single cell suspensions as described above. Approximately 1 million cells were transferred to KP.SIY tumor bearing mice seven days post tumor inoculation. Recipient animals were euthanized and analyzed three days post 2C T cell transfer. For transfers into secondary recipients, 2C T cells were transferred into primary recipients as described above. Three days post adoptive transfer, the tumor-draining iLN or spleen of recipient mice were isolated and CD8^+^ T cells were enriched with the Miltenyi CD8^+^ T cell isolation kit. Congenically marked 2C T cells were then isolated from the CD8^+^ T cells using fluorescence activated cells sorting and 50,000 2C T cells were transferred intravenously into secondary recipient C57BL/6 mice bearing day 7 flank KP.SIY tumors. Secondary recipients received ICB one and three days after adoptive transfer of in vivo primed 2C T cells and were analyzed seven days after adoptive transfer of in vivo primed 2C T cells.

### In vivo antigen delivery

C57BL/6 splenocytes were isolated as described above, then ACK lysed, washed, and pulsed in complete medium for 1 hour at 37 degrees Celsius with 0.2 μM SIY peptide. SIY-pulsed splenocytes were washed and injected intravenously into KP.SIY tumor-bearing mice on days 4, 6, and 9 post tumor inoculation. During this experiment, some mice were also treated with ICB on days 7 and 10 post-tumor inoculation. Mice were analyzed on day 14 post-tumor inoculation.

### Single-cell RNA sequencing with Seq-Well

Sorted cells were then processed for scRNA-seq using the Seq-Well platform with second strand chemistry, as previously described (52, 53). Whole transcriptome libraries were barcoded and amplified using the Nextera XT kit (Illumina) and were sequenced on a Novaseq 6000 (Illumina). Hashtag oligo libraries were amplified as described previously and were sequenced on a Nextseq 550(54).

### Processing of cell hashing data

Cell hashing data was aligned to HTO barcodes using CITE-seq-Count v1.4.2. First, cells receiving fewer than five total HTO counts were classified as negatives. To establish thresholds for positivity for each HTO barcode, we first performed centered log-ratio normalization of the HTO matrix and then performed k-medoids clustering with k=5 (one for each HTO). This produced consistently five clusters, each dominated by one of the 5 barcodes. For each cluster, we first identified the HTO barcode that was dominant in that cluster. We then considered the threshold to be the lowest value for that HTO barcode among the cells classified in that cluster. To account for the scenario in which this value was substantially lower than the rest of the values in the cluster, we used Grubbs’ test to determine whether this threshold was statistically an outlier relative to the rest of the cluster. If the lower bound was determined to be an outlier at p=0.05, it was removed from the cluster, and the next lowest value was used as the new threshold. This procedure was iteratively applied until the lowest value in the cluster was no longer considered an outlier at p=0.05. Cells were then determined to be “positive” or “negative” for each HTO barcode based on these thresholds. Cells that were positive for multiple HTOs or were negative for all HTOs were excluded from downstream analysis. To account for differences in sequencing depth between samples, these steps were performed separately for each Seq-Well array that was processed. Thresholds calculated for each sample were manually inspected and adjusted if necessary. Cells marked as “doublets” or “negatives” by this procedure were excluded from downstream analysis.

### Processing of single-cell RNA sequencing data

Raw read processing of scRNA-seq reads was performed as previously described(55). Briefly, reads were aligned to the mm10 reference genome and collapsed by cell barcode and unique molecular identifier (UMI). Then, cells with less than 300 unique genes detected or with greater than 25% mitochondrial gene counts and genes detected in fewer than 5 cells were filtered out. Cell cycle scores for individual cells were computed using CellCycleScoring function in Seurat. Data was then integrated by batch using Seurat v4.1.1(55). The ScaleData function was then used to regress out the number of RNA features in each cell, as well as S and G2/M cell cycle scores and fraction of mitochondrial gene expression. The number of principal components used for visualization was determined by examination of the elbow plot, and two-dimensional embeddings were generated using uniform manifold approximation and projection (UMAP). Clusters were determined using Louvain clustering, as implemented in the FindClusters function in Seurat. DEG analysis was performed for each cluster and between indicated cell populations using the FindMarkers function. Data was iteratively reclustered to remove clusters with gene expression consistent with naïve T cells, monocytes, and NK cells. Label transfer of cluster labels onto proliferating cell populations was performed using the FindTransferAnchors and TransferData functions in Seurat (55).

### Reanalysis of published single-cell datasets

Data from Giles et al. and Daniel et al. were obtained from GSE199565 and GSE188670. To generate signatures, we determined the top 50 differentially expressed genes. For Giles et al., data from day 8, day 15, and day 30 of clone 13 LCMV infection were used. For Daniel et al. GP33 tetramer-positive cells from the spleen of day 21 of LCMV infection were used. Cell scores were then determined using these gene lists and the AddModuleScore function in Seurat (55). Data from Zheng et al was downloaded from Zenodo. Four component datasets that contained transcript expression and T cell receptor data from both tumor tissue and matched peripheral blood (CRC.LeiZhang2018, HCC.ChunhongZheng2017, NSCLC.XinyiGuo2018, and CHOL.thisStudy) was aggregated and integrated by patient. Enrichment of mouse exhausted signatures was performed using the AddModuleScore function in Seurat. TCR data was obtained from the file tcr.zhangLab.comb.flt.rds. A clonotype was considered “tumor-trafficking” or “blood-related” if it contained at least one cell in either the tumor or the peripheral blood, respectively. Two patients that exhibited no clonotypes matched between peripheral blood and tumor were excluded from this analysis.

### Paired single-cell TCR sequencing and analysis

Paired TCR sequencing and read alignment was performed as previously described (56). Briefly, whole transcriptome amplification product from each single-cell library was enriched for TCR transcripts using biotinylated *Tcrb* and *Tcra* probes and magnetic streptavidin beads. The enrichment product was further amplified using V-region primers and Nextera sequencing handles, and the resulting libraries were sequenced on an Illumina Novaseq 6000 or Nextseq 550. Processing of raw sequencing reads was performed using the Immcantation software suite (57, 58). First, the FilterSeq.py function was used to remove reads with an average quality score less than 25. Then, reads were aggregated by cell barcode and UMI, and UMI with under 10 reads were discarded ClusterSets.py was used to divide sequences for each UMI into sets of similar sequences. Only sets of sequences that comprised greater than 70% of the sequences obtained for that UMI were considered further. Consensus sequences for each UMI were determined using the BuildConsensus.py function. Consensus sequences were then mapped against TCRV and TCRJ IMGT references sequences with IgBlast (59). Sequences for which a CDR3 sequence could not be unambiguously determined were discarded. UMI for consensus sequences were corrected using a directional UMI collapse, as implemented in UMI-tools. TCR sequences were then mapped to single cell transcriptomes by matching cell barcodes. If multiple *Tcra* or *Tcrb* sequences were detected for a single cell barcode, then the corresponding sequence with the highest number of UMI and raw reads was retained. To define clonotypes of cells, we first segregated cells by mouse and unique *Tcrb* CDR3 junction nucleotide sequences. For each unique combination of mouse and CDR3β junction, we determined the most common TCRα sequence in cells with paired TCR recovery. We then imputed missing beta chains from cells with recovery of only alpha chain by matching to these combinations of mouse, beta chain, and alpha chain.

