## Supplement Figures for "Differentiation and Expansion of Tumor-Infiltrating T Cell Clonotypes Occurs in the Spleen Following Immune Checkpoint Blockade"

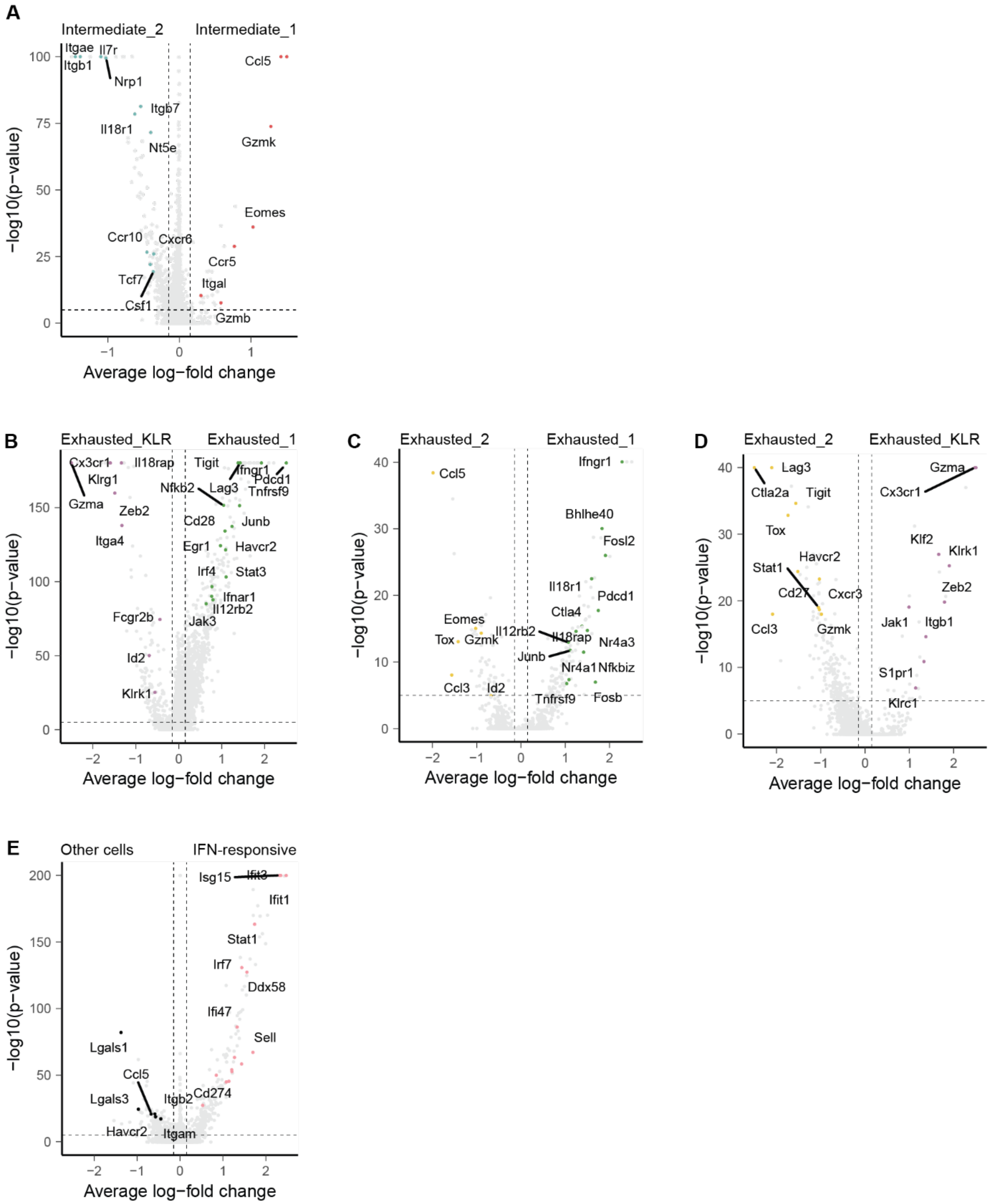


**Supplemental Figure 1**. A) Volcano plot of differentially expressed transcripts between Intermediate_1 and Intermediate_2 cell phenotypes. B) Volcano plot of differentially expressed transcripts between Exhausted_1 and Exhausted_KLR cell phenotypes. C) Volcano plot of differentially expressed transcripts between Exhausted_1 and Exhausted_2 cell phenotypes. D) Volcano plot of differentially expressed transcripts between Exhausted_KLR and Exhausted_2 cell phenotypes. E) Volcano plot of differentially expressed transcripts between IFN-responsive cells and all other SIY-reactive cells.

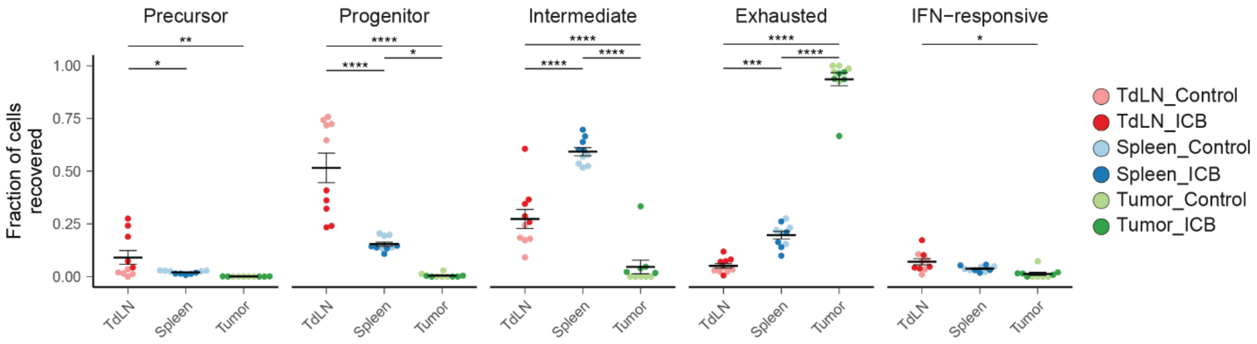


**Supplemental Figure 2**. Frequency of pre-exhausted, progenitor, intermediate, exhasuted, and IFN-responsive phenotypes in the TdLN, spleen, and tumor. P-values are calculated using one-way ANOVA followed by a Tukey honestly significant difference (HSD) test. Error bars show mean +/- standard error of the mean.

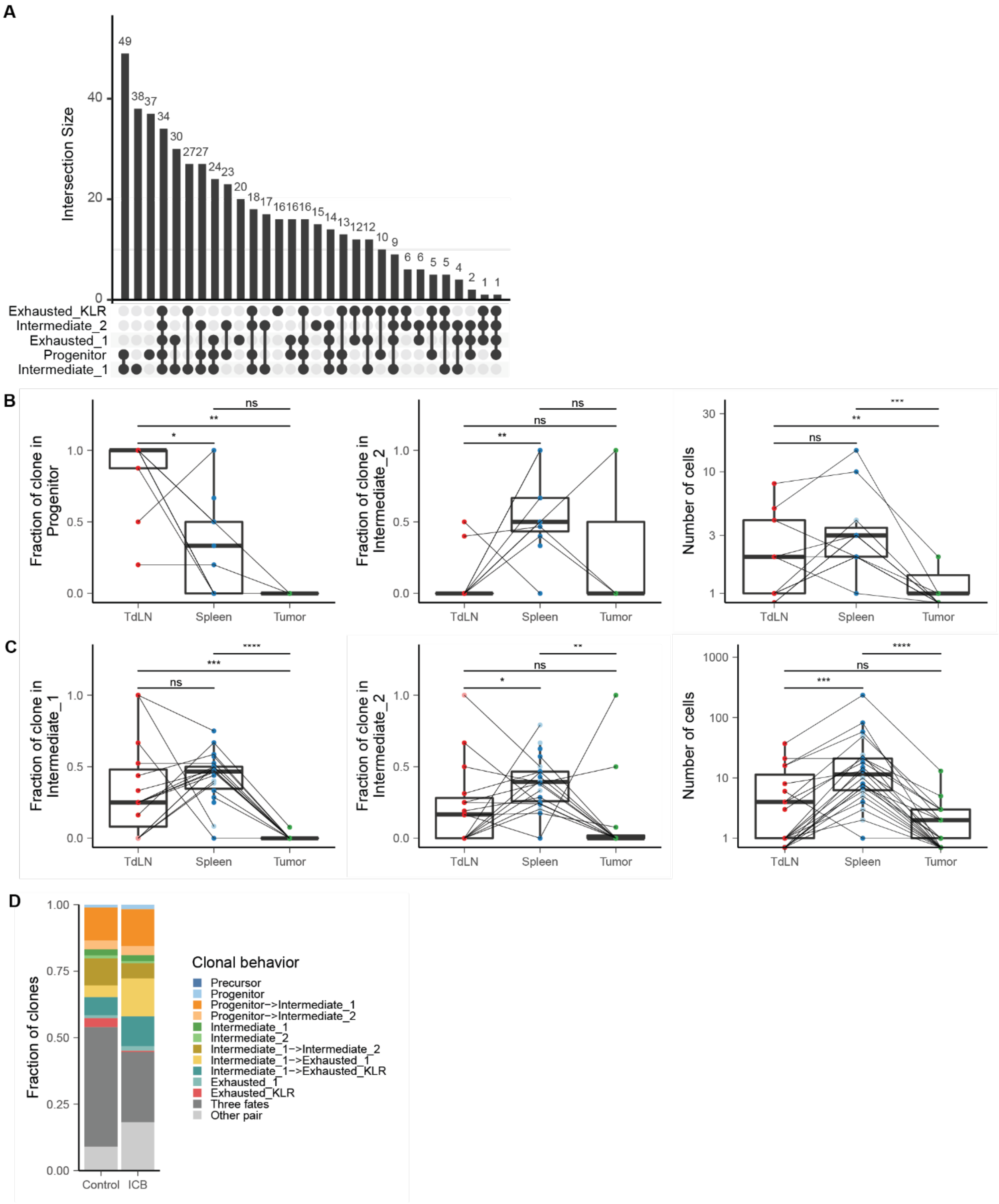


**Supplemental Figure 3**. A) Upset plot depicting patterns of phenotype sharing within individual clonotypes. All clonotypes shown have clonal sizes greater than one. B) Tissue-site distribution of progenitor and intermediate_2 phenotypes and clonal sizes in progenitor->intermrediate_2 clones. P-values are calculated with a two-sided Wilcoxon rank-sum test and are adjusted with Bonferroni correction. C) Tissue-site distribution of intermediate_1 and intermediate_2 phenotypes and clonal sizes in intermediate_1->intermrediate_2 clones. P-values are calculated with a two-sided Wilcoxon rank-sum test and are adjusted with Bonferroni correction. D) Stacked bar plot of clonal behaviors in control and ICB-treated mice, weighted by number of clonotypes. Only clonotypes with a clonal size > 2 are shown.

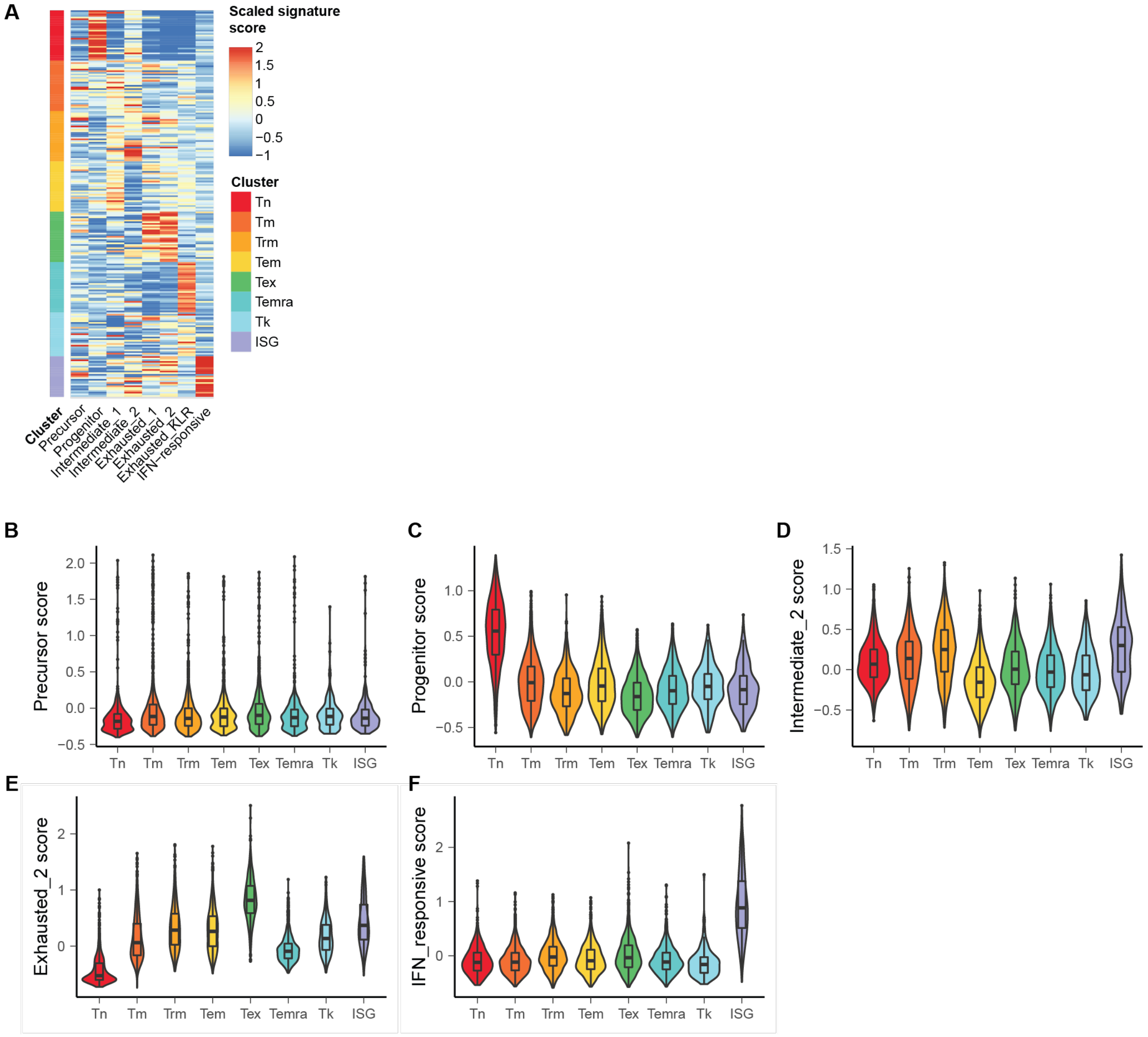


**Supplemental Figure 4.** A) Heat map of scaled signature scores expressed by phenotypes defined by Zheng et al. Each row represents the average expression of each signature in cells from a single patient. B-F) Signature scores for precursor (B), progenitor (C), intermediate_2 (D), exhausted_2 (E), and IFN-responsive (F) phenotypes..
